# Gene Cascade Finder: A tool for identification of gene cascades and its application in *Caenorhabditis elegans*

**DOI:** 10.1101/593400

**Authors:** Yusuke Nomoto, Yukihiro Kubota, Kota Kasahara, Aimi Tomita, Takehiro Oshime, Hiroki Yamashita, Muhamad Fahmi, Masahiro Ito

**Affiliations:** Advanced Life Sciences Program, Graduate School of Life Sciences, Ritsumeikan University, Kusatsu, Shiga, Japan; Department of Bioinformatics, College of Life Sciences, Ritsumeikan University, Kusatsu, Shiga, Japan

## Abstract

Obtaining a comprehensive understanding of the gene regulatory networks, or gene cascades, involved in cell fate determination and cell lineage segregation in *Caenorhabditis elegans* is a long-standing challenge. Although RNA-sequencing (RNA-Seq) is a promising technique to resolve these questions, the bioinformatics tools to identify associated gene cascades from RNA-Seq data remain inadequate. To overcome these limitations, we developed Gene Cascade Finder (GCF) as a novel tool for building gene cascades by comparison of mutant and wild-type RNA-Seq data along with integrated information of protein-protein interactions, expression timing, and domains. Application of GCF to RNA-Seq data confirmed that SPN-4 and MEX-3 regulate the canonical Wnt pathway during embryonic development. Moreover, *lin-35, hsp-3*, and *gpa-12* were found to be involved in MEX-1-dependent neurogenesis, and MEX-3 was found to control the gene cascade promoting neurogenesis through *lin-35* and *apl-1*. Thus, GCF could be a useful tool for building gene cascades from RNA-Seq data.

## Introduction

Spatially and temporally regulated gene expression is essential to precisely modulate cellular behaviors during development in multicellular organisms. Elucidating gene cascades during early embryonic development may improve our understanding of mechanisms of cell fate determination and lineage segregation [1-3]. The nematode *Caenorhabditis elegans*, a model organism of development research, comprises 959 cells in adult hermaphrodites with robustness and reproducibility of the cell lineage [3]. Additionally, over 80% of the *C. elegans* proteome shows homology with human proteins [4], providing a particularly valuable model organism for studies of the developmental system.

PAR proteins, which are expressed immediately after fertilization, are associated with formation of the anterior-posterior polarity of P0 cells and control the localization of polarity mediators such as SPN-4, MEX-1, and MEX-3 in *C. elegans* [5]. Aberrant expression of the genes encoding these proteins affects cellular and developmental regulation, leading to embryonic lethality in early embryogenesis [6-9]. Specifically, SPN-4 is localized in all blastomeres at the four-cell stage and plays essential roles in axial rotation [8, 10-12]; MEX-1 is expressed in the P1 blastomere, and loss of its function leads to excessive muscle formation [7, 13]; and MEX-3 is expressed in the AB blastomere, and causes excessive muscle formation and hatching failure in mutants [14-16]. Although polarity mediators regulate protein synthesis by binding to the 3′-untranslated region of the target mRNA, it is difficult to directly identify their associated gene cascades.

Conventional genetic and molecular biological approaches have focused on the target gene to be identified and have clarified functions and identified related genes, representing a bottom-up approach. Both functional analysis of individual genes and comprehensive analysis of the genome are indispensable for identification of gene cascades. After determination of the whole genome sequence of *C. elegans* in 1998, genome-wide analysis via a top-down approach was made possible [17], representing the beginning of the post-genome sequencing era. Transcriptomics via DNA microarray analysis [18], proteomics via mass spectrometry analysis [19], and phenomenon analysis by RNA interference [20] have been extensively reported. Furthermore, several methods for comprehensive analyses have been developed, including protein-protein interaction analysis using a yeast two-hybrid system and phage display [21, 22] and multiple mutation analysis using knockdown mutants. At the same time, the WormBase database was constructed to integrate the vast quantities of data obtained from these genome-wide analyses [23]. Accordingly, the development of new technologies and methodologies has enabled the accumulation of detailed genome-wide data.

Next-generation sequencing (NGS) has now replaced conventional Sanger sequencing [24]. Conventional Sanger sequencing can simultaneously analyze 8–96 sequencing reactions, whereas NGS can simultaneously run millions to billions of sequencing reactions in parallel. This technique can dramatically and quickly determine the gene sequences in organisms whose whole-genome sequences have already been determined. Even at the laboratory level, genomic sequencing results can be produced in only a few days, enabling researchers to obtain genome-wide information rapidly. Furthermore, RNA sequencing (RNA-Seq) has recently been developed to measure gene expression levels by counting the number of sequence reads obtained from converting RNA into cDNA [25]. Existing RNA-Seq data analysis tools include RSeQC [26], which measures the quality control of the obtained data; Cufflinks [27], which involves genome mapping; and IsoEM [28], which identifies isoforms within a dataset. These tools can be used to identify gene expression variations from RNA-Seq data. TopGO (Alexa et al., 2006) is an analytical tool used to identify gene function based on RNA-Seq data and can confirm the functions of genes with varying expression levels. In addition, Cascade R was established to identify the gene cascade of a query gene [29]. Cascade R constructs an intergenic network of knockout genes from the results of DNA microarray analysis. However, it requires multiple timeline datasets from microarray analyses.

Genes, especially those expressed in early embryogenesis, function in chronological order rather than having only a single function, and genes responsible for functional expression often exert their effects at the bottom of gene cascades. STRING [30], BIOGRID [31], and WormBase [23] are databases of protein-protein interactions and the gene-dependent regulation of transcription and translation. In order to predict genetic cascades from these databases, researchers currently must perform separate analyses. Moreover, although RNA-Seq can be used to easily acquire large amounts of data via a semi-automatic process, the subsequent analysis must be performed manually and is therefore quite time-consuming. Therefore, the data acquisition capacity currently exceeds the data analysis capacity. Accordingly, automation of the analysis using bioinformatics tools is an important research subject.

In this study, we performed RNA-Seq analysis of the polarity mediator mutants *spn-4, mex-1*, and *mex-3* in C. elegans. Next, we developed a novel tool, Gene Cascade Finder (GCF), to extract genes with a high probability of being directly or indirectly regulated by these polarity mediators. Finally, the gene cascade and its validity were examined.

## Methods

### Strains

*C. elegans* N2, *mex-1* (or286), and *mex-3* (eu149) strains, and *Escherichia coli* OP50 strain were provided by the Caenorhabditis Genetics Center (https://cgc.umn.edu/), and the *C. elegans spn-4* (tm291) strain was provided by National BioResource Project [32].

### Culture of *C. elegans* and synchronization at the early embryo stage

All strains except for *mex-1* (or286) were cultured on nematode growth medium agar coated with *E. coli* OP50 at 20°C. Because *mex-1* (or286) strain is a temperature-sensitive mutant strain, it was cultured at 15°C to strengthen its phenotype [13]. Furthermore, all strains were transferred to S-Basal solution inoculated with *E. coli* OP50 at 20°C for large culture. To obtain early embryos from the culture medium, when *C. elegans* adults had only 3–5 eggs, they were synchronized using an alkaline bleaching method, and the early embryos were recovered [33]. These *C. elegans* early embryos were used as the samples for RNA-Seq analysis.

### RNA-Seq analysis

The mRNAs of the synchronized *C. elegans* early embryos were purified using RNeasy Minikit (Qiagen NV, Venlo, the Netherlands). Purified mRNAs were reverse-transcribed into cDNAs, amplified by polymerase chain reaction, and fragmented using a TruSeq RNA Sample Prep Kit (Illumina, Inc.). The amplified cDNAs were sequenced using Hi-Seq2000 (Illumina, Inc.) and the sequenced cDNAs were mapped to the *C. elegans* genome sequence and counted according to WormBase (WS190) [23] using DNAnexus. Using this procedure, the mRNA expression levels were obtained as reads per kilobase of exon per million mapped reads (RPKM) [34]. The gene name and RPKM values of wild-type and mutant genes were filed for input data in GCF (S1 Table).

### Comparative quantitative gene expression analysis

Expression levels of gene *i* in the wild-type and mutant were defined as x*i* and y*i*, respectively, and the change rates in these gene expression levels (R*i*) were determined as shown in Equation 1. Because the data obtained by RNA-Seq analysis had a non-normal distribution, the data were subjected to non-parametric tests using Equations 2a and 2b.

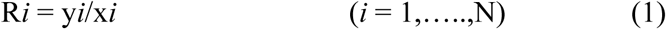

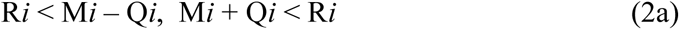

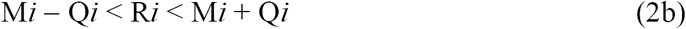

Where N is the number of genes, and M*i* and Q*i* are the median and quartile deviation, respectively. Genes that satisfied the condition of Equation 2a were assumed to show expression level fluctuations, and genes satisfying the condition of Equation 2b were assumed to not show expression level fluctuations.

### Dataset for the software

Information on the expression timing and interactions of all genes in *C. elegans* was extracted from WormBase (Version 256) [23] using the application programming interface. The total transcription factors of *C. elegans* were acquired from the gene ontology database Amigo 2 (http://amigo.geneontology.org/amigo) [35] using the keyword search “GO: 0006351”. Furthermore, all gene IDs, protein IDs, and domain information from Pfam (https://pfam.xfam.org/) [36] in *C. elegans* required for functional analysis were extracted from UniProt [37].

### Direct target prediction by GCF

GCF was developed by the algorithm shown in Fig. 1. First, the candidate genes of the target of the query gene were found from transcription factors and genes with no gene expression level fluctuations, as calculated by Equation 2b, with the same cellular localization and phenotype. Query genes bind to target mRNAs to regulate their translation. Thus, the mRNA expression levels of the target genes showed no changes. In addition, the target genes needed to be expressed with the same timing as the query gene because the query gene is directly bound to the target mRNA. Furthermore, the gene cascade of the target genes was mostly consistent with the query gene cascade, suggesting that the target gene may have the same phenotype as the query gene. Therefore, to expand the gene cascade, the target gene should be a transcription factor with downstream genes.

**Fig. 1.**
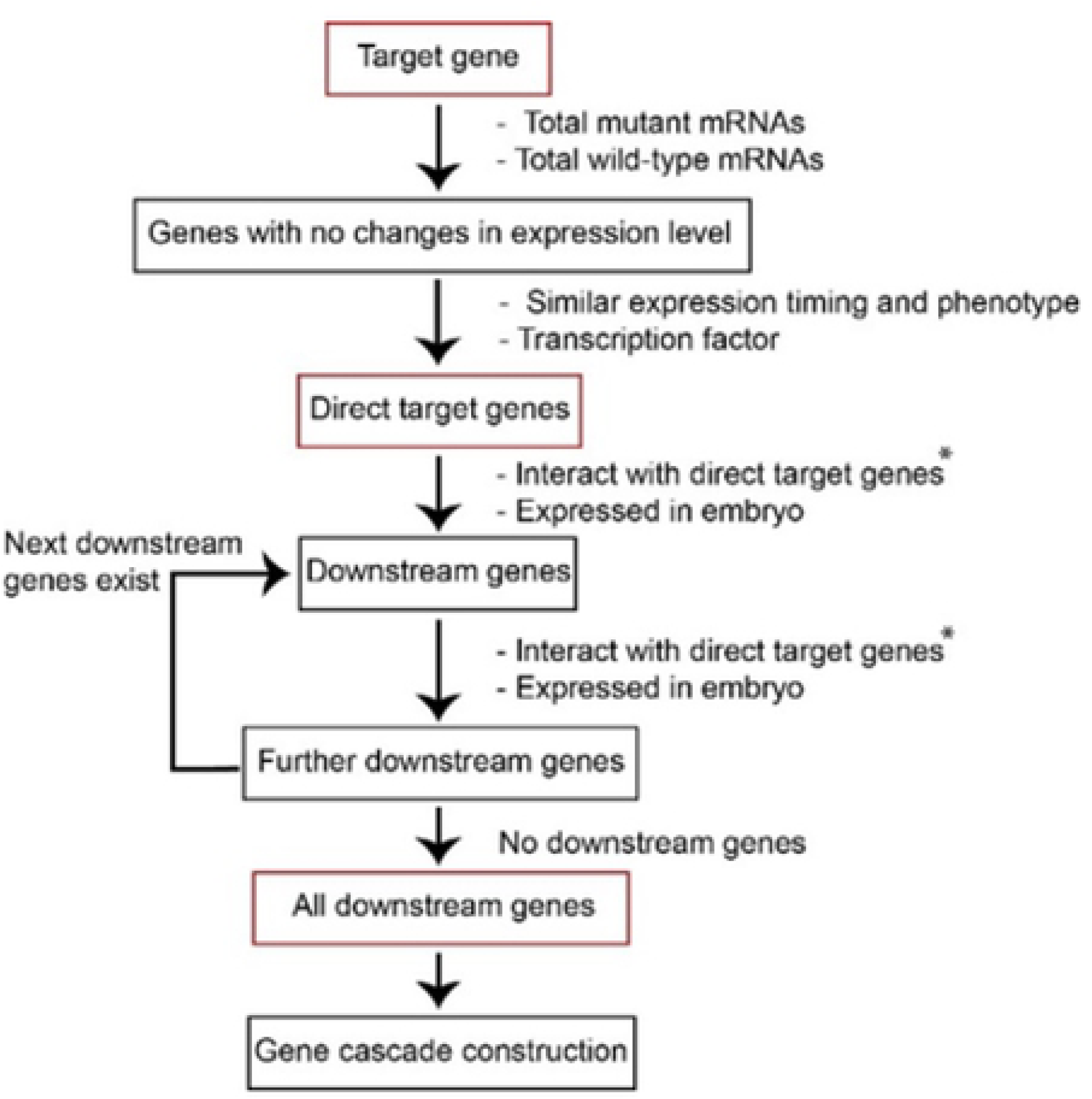
Schematic for prediction of the gene cascade. Application protocol for GCF. Genes in the predicted cascade are indicated by red frames. To identify the entire gene network, we repeatedly identified the downstream genes. The genes surrounded by black frames were required to identify the genes surrounded by red fames. The information of the labels with asterisks was extracted from WormBase.

### Downstream gene identification by GCF

The search for downstream genes was carried out as follows. First, the genes from transcription or protein-protein interactions were extracted as downstream gene candidates of the target gene. Second, the expression timing of the candidate genes was checked. Only candidate genes noted as being expressed in early embryos or in embryos in WormBase were defined as downstream genes of the query’s target genes. These first two steps were then repeated to obtain the next downstream genes. Finally, the procedure was repeated until there were no genes left to be extracted to obtain the final gene. Lastly, GCF output the cascade data (S2–S4 Tables). An example showing input obtained from output data from GCF to Cytoscape [38] is provided in Fig. 2.

**Fig. 2.**
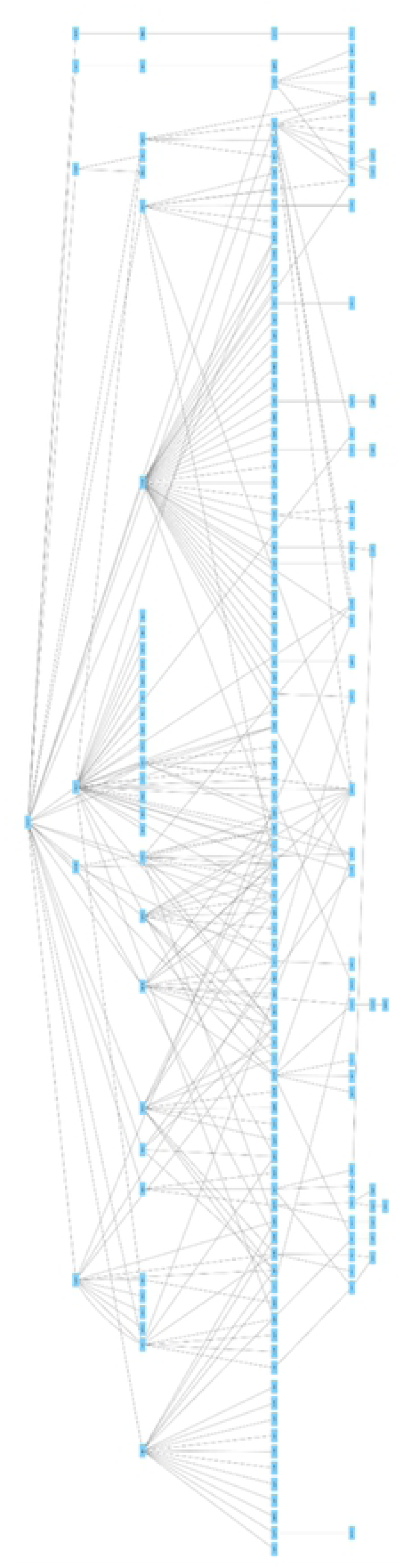
Representative example of the graphic output from GCF. Graphical examples from the GCF software were further processed using Cytoscape, allowing for identification of an output without adding a new input.

### Specific domain search from the constructed gene cascade

Each direct target gene was rooted, and the functions of their bottom genes were investigated using Pfam [36] in UniProt [37]. The P-values of the domains from the bottom gene products were evaluated using the same formula for Gene Ontology in Panther [39]. If the transcription-related domain was extracted from a cascade, the cascade was no longer considered since it would be functioning only after the early embryo stage.

## Results and Discussion

### Development of Gene Cascade Finder

The programming language Ruby was used to construct Gene Cascade Finder (GCF). The web interface of GCF was written in Python. The input data for GCF were data from the wild-type and mutant strains as shown in S1 Table. The output data from GCF were tables of discovered gene cascades (S2–S4 Tables), the data input into Cytoscape (S5 Table), and gene cascade-specific domains and their gene cascades (S6 Table). GCF is available at http://www.gcf.sk.ritsumei.ac.jp

### Analysis of mRNA expression by comparative RNA-Seq

To explore polarity mediator-dependent mechanisms, the effects of deficiencies in polarity mediators were analyzed by performing RNA-Seq analysis in early embryos. From the results of comparative RNA-Seq of the wild-type strain and the *spn-4, mex-1*, and *mex-3* mutant strains, 15,288, 15,265, and 15,005 genes were identified, respectively (S7 Table). In these gene groups, expression level fluctuations were calculated by examining the median ± quartile deviations. From this analysis, 6,417 genes distributed at −0.65 < log2 (RNA expression level ratio) < 0.69 in the *spn-4* gene, 6,456 genes distributed at −0.74 < log2 (RNA expression level ratio) < 0.82 in the *mex-3* gene, and 6,491 genes distributed at −0.82 < log2 (RNA expression level ratio) < 1.10 in the *mex-1* gene were defined as genes showing no expression level variations (S8 Table).

### Gene cascade prediction using Gene Cascade Finder

As shown in S1 Table, gene cascade prediction was performed by inputting data obtained by comparative RNA-Seq into GCF. GCF can predict gene cascades by continuously integrating the results from RNA-Seq along with data on gene expression and intermolecular interactions from WormBase. In total, 127, 180, and 226 gene cascades were predicted from 6,418, 6,457, and 8,513 genes from the comparative analysis of the *spn-4, mex-1*, and *mex-3* mutant strains, respectively (Fig. 3 and S6–S8 Tables).

**Fig. 3.**
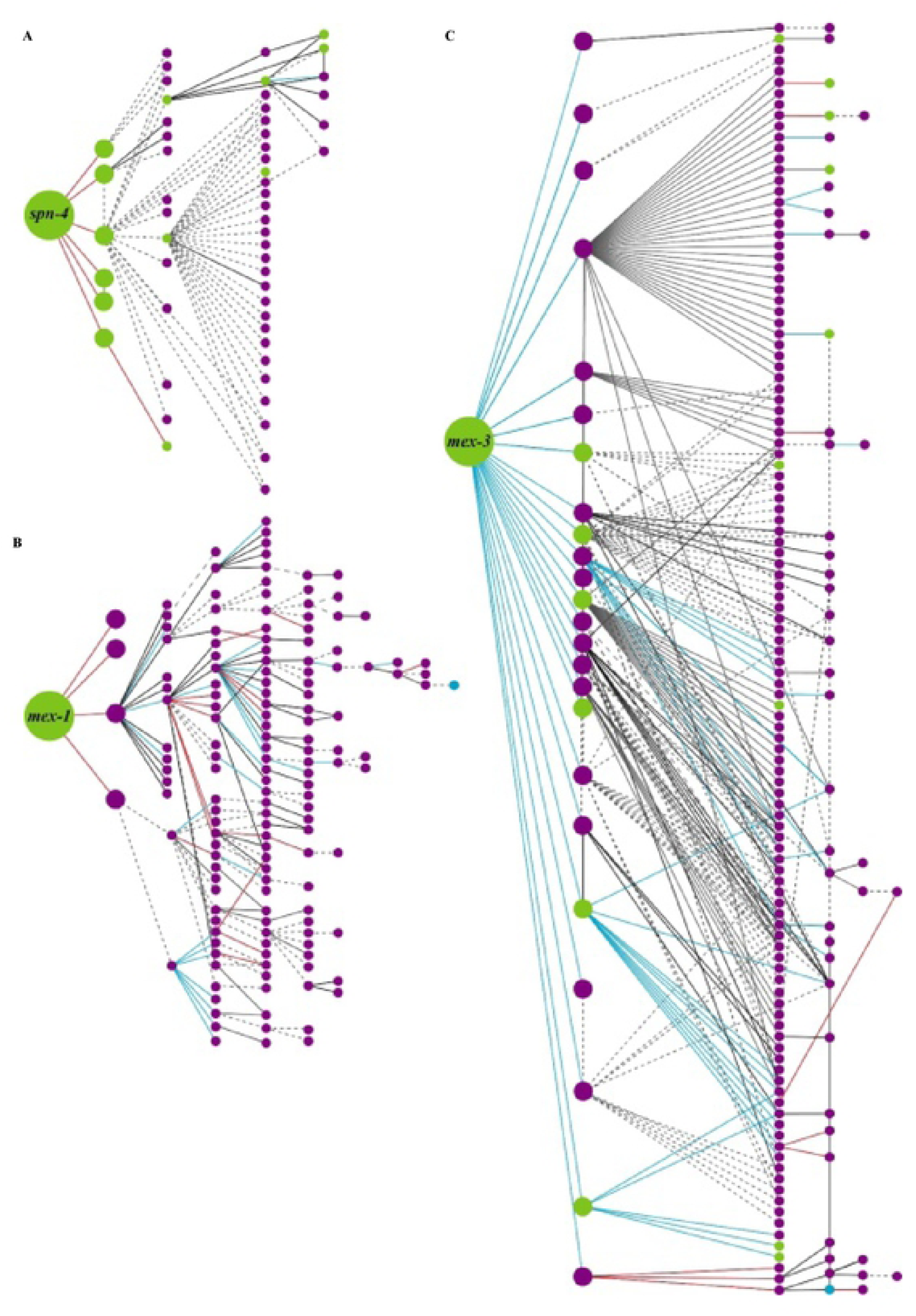
Schematic illustrations of polarity mediator-dependent gene cascades during *C. elegans* embryogenesis. Rendering of each gene cascade was performed using Cytoscape. Nodes indicate each gene in the cascade. Edges indicate the interactions between two genes. Large and intermediate nodes indicate the query genes and the direct target of the query genes, respectively. Other nodes indicate downstream genes. Green nodes indicate genes that are expressed during the early embryonic stage. Purple nodes indicate presumptive early embryonic genes. Red, blue, and black edges indicate positive regulation, negative regulation, and genetic interactions, respectively. Dotted lines indicate protein-protein interactions. (A) *spn-4*-mediated gene cascade. (B) *mex-1*-mediated gene cascade. (C) *mex-3*-mediated gene cascade.

### Extraction of gene cascade-specific domains using Gene Cascade Finder

The genes and domains located at the bottom of the gene cascade were extracted for functional analysis of the predicted gene cascade (S9–S11 Tables). Overall, 53, 146, and 143 genes with 32, 34, and 54 specific domains as the bottom gene were extracted from the gene cascades in the *spn-4, mex-1,* and *mex-3* mutants, respectively.

### Domain analysis of *spn-4, mex-1*, and *mex-3* cascades

To predict the functions of the gene cascades, we focused on the functions of genes localized at the bottom of the gene cascade by analyzing the domains of the gene products using the Pfam protein family database. The functions of the 53 SPN-4-mediated genes were obtained as bottom genes to calculate the functional trends in the *spn-4* cascade. Within the 53 bottom genes, 32 domains were classified based on information from the Pfam database (Table 1) (Bateman et al., 2004). When we calculated the numbers of these genes, transcription and signal transduction were obtained at high frequency.

**Table 1.** Scores of the functional characterization of *spn-1-, mex-1-*, and *mex-3*-mediated gene cascades by domain analysis. Properties of the gene product of the bottom genes were calculated by domain analysis. The sum of the characteristic features of each cascade (P < 0.05) was then calculated. Domains with a score less than 1 were classified as “Others”.

|  | *spn-4* | *mex-1* | *mex-3* |
| --- | --- | --- | --- |
| Transcription | 25 | 6 | 4 |
| Signal transduction | 5 | 3 | 12 |
| Development | - | 2 | 7 |
| Cell cycle | - | 3 | 5 |
| Cell division | - | - | - |
| DNA replication | - | 2 | 3 |
| Transport | - | 3 | - |
| Others | 2 | 12 | 18 |
| Unknown | - | 3 | 4 |

Similarly, in the gene cascade of 146 *mex-1*-mediated genes, 27 domains were classified and obtained from the domain analyses to have functions in early embryonic development, cell division, transcription, DNA replication, and signal transduction (Table 1). In contrast, in the gene cascades of the 143 *mex-3*-mediated genes, 54 domains were classified and obtained from the domain analyses to have functions in development, cell cycle, transcription, and signal transduction (Table 1).

### Evaluation of GCF by assessment of gene cascades in the canonical Wnt signaling pathway

Next, we focused on genes involved in SPN-4-mediated signal transduction (Fig. 4A). First, we found that MOM-2, a nematode homolog of the Wnt ligand, is involved in the signal transduction cascade (Table 1). Since both SPN-4 and MOM-2 were previously reported to regulate EMS cell lineage formation and spindle orientation [11, 40], we hypothesized that the SPN-4/MOM-2 gene cascade may have an essential role in early embryogenesis and may regulate the Wnt signaling pathway. Moreover, because OMA-1, MEX-1, and PIE-1, which are known to be essential for MOM-2 expression in embryonic development [40], were also identified in this pathway, the GCF-mediated gene cascade prediction was assumed to be accurate.

**Fig. 4.**
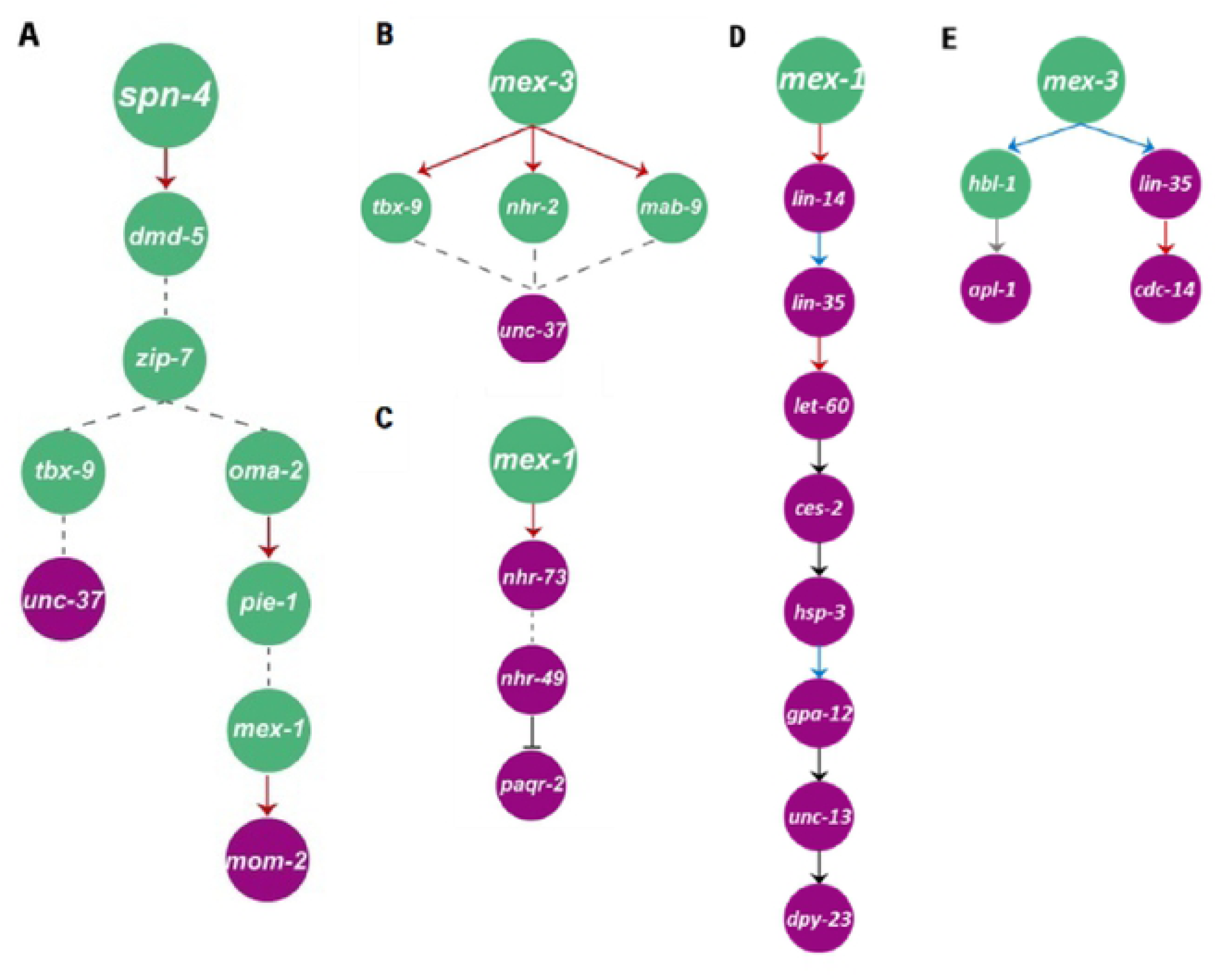
Typical examples of SPN-4, MEX-1, and MEX-3-dependent gene cascades. (A) SPN-4-mediated gene cascade regulates Wnt signaling. (B) MEX-3-mediated gene cascade negatively regulates Wnt signaling. (C) MEX-1-mediated gene cascade regulates endoplasmic reticulum associated degradation (ERAD) of folding-deficient proteins. (D) MEX-1-mediated gene cascade regulates neuronal development. (E) MEX-3-mediated gene cascade regulates neuronal development.

Similarly, *unc-37*, which encodes a Groucho/TLE homolog that suppresses Wnt signaling, was found as the “bottom gene” in the *spn-4* and *mex-3* cascades (Fig. 4A, B). Thus, we propose that GCF-mediated gene cascade prediction may be useful for identification of gene cascades involved in *C. elegans* embryonic development.

### Examples of the application of GCF for prediction of new biological functions involved in a gene cascade

#### Endoplasmic reticulum (ER) stress response pathway in a MEX-1-mediated gene cascade

Because cell division and DNA replication are regulated by a MEX-1-dependent cascade, we further focused on this signal transduction cascade. In the gene cascade related to signal transduction, *paqr-2*, which encodes an adiponectin receptor, was isolated (Fig. 4C). Interestingly, a stress response pathway is known to regulate stress responses in both mouse and *C. elegans* embryogenesis [40, 41]. Thus, an evolutionarily conserved gene cascade against environmental stress may be identified using GCF. Moreover, because adiponectin receptor regulates insulin signaling [42], it is likely that MEX-1/PAQR-2-mediated gene cascades may be involved in ER stress tolerance signaling within or parallel to the insulin signaling pathway during embryogenesis.

#### *lin-35, hsp-3,* and *gpa-12* in a MEX-1/DPY-23-mediated gene cascade in neuronal development

In a MEX-1/DPY-23-mediated gene cascade, the *mex-1, lin-14, let-60, ces-2, unc-13*, and *dpy-23* mutants were shown to exhibit specific phenotypes in neuronal development (Fig. 4D) [12, 43-45]. Thus, six of the nine genes of this gene cascade were shown to have essential roles in neuronal development, indicating that MEX-1/DPY-23-mediated gene cascades may regulate neuronal function. Although their roles in neuronal development have not yet been investigated, locomotion defects have been reported in *hsp-3* and *gpa-12* mutants [46, 47]. Similarly, the *lin-35* (n745) mutation has been shown to enhance the neuronal phenotype of the neuronal regulator genes *dpy-13* and *unc-104* [48]. Thus, *lin-35, hsp-3*, and *gpa-12* may be involved in a DPY-23-mediated gene cascade in neuronal development in embryos [45]. However, further studies are required to examine this possibility.

#### MEX-3/APL-1-mediated neuronal patterning and MEX-3/CDC-14-mediated cell fate determination in the MEX-3-mediated gene cascade

Because MEX-3 is specifically expressed in AB cells at the four-cell stage, spatiotemporal-regulated synaptic formation defects in *hbl-1* mutants and *apl-1*-dependent embryonic neuronal patterning may be elucidated by identifying MEX-3/APL-1-mediated gene cascades (Fig. 4E) [49-51]. In parallel, when we focused on the MEX-3/CDC-14-mediated gene cascade (Fig. 4E), CDC-14B, a zebrafish homolog of CDC-14, was shown to be involved in formation of the cilium in sensory neurons [50]. Because sensory neurons have cilia in *C. elegans* [52], CDC-14 may be involved in an evolutionarily conserved signaling pathway. Similarly, the *lin-35* (n745) mutation was shown to enhance the neuronal phenotype of neuronal regulator genes [48]. Thus, *lin-35* may be involved in a MEX-3/CDC-14-mediated gene cascade in sensory neuron development. Accordingly, our findings suggested that GCF may be useful for predicting the comprehensive functions of query genes and for identification of new genes involved in known gene cascades.

## Conclusion

In this study, we created a software program called GCF, which could comprehensively identify genes downstream of the query genes by integrating RNA-Seq data and previously characterized data from WormBase. Using GCF, we analyzed gene cascades of the polarity mediator proteins SPN-4, MEX-1, and MEX-3, and identified 127, 180, and 226 putative gene cascades, respectively. By analyzing the functions of these gene cascades, we confirmed that SPN-4 and MEX-3 regulate the canonical Wnt pathway during embryonic development. Furthermore, we found that the ER stress response and motor neuron development are regulated by MEX-1-dependent cascades, and that neural development is regulated by MEX-3-dependent cascades. Although we used GCF only to evaluate SPN-4, MEX-1, and MEX-3 functions in this study, the method is applicable for other translation or transcription factors involved in early embryogenesis. In addition, GCF provides a general method for predicting the functions of genes involved in a gene cascade during *C. elegans* embryonic development. Taken together, we propose that our strategy using the GCF tool offers a reliable approach for comprehensively identifying networks of embryo-specific gene cascades in *C. elegans*. Importantly, GCF can also be applied to humans and other model organisms such as mice and *Drosophila*.

In the future, by expanding the algorithm to fit the cell lineage-segregation of *C. elegans* [3], we will be able to predict the precise gene cascades reflecting four-dimensional (spatial and temporal) regulation [53]. Combinational analysis of GCF and molecular biology techniques such as RNA-pull down assays, fluorescent *in situ* hybridization, and phenotypic characterization of the mutants may be required to build a more reliable regulatory network for these gene cascades [54, 55].

## Acknowledgements

We would like to thank Dr. Hisao Kojima, Mr. Takahiro Nakamura, and Mr. Yuuto Ohnishi for their support and helpful comments, and Mr. Marori Yoshioka and Dr. Takuya Takahashi for fruitful discussions.

## References

1. English J, Pearson G, Wilsbacher J, Swantek J, Karandikar M, Xu S, et al. New insights into the control of MAP kinase pathways. Experimental cell research. (1999);253(1):255–70. Epub 1999/12/02. doi:10.1006/excr.1999.4687. PubMed PMID:10579927.

2. Maduro MF, Rothman JH. Making worm guts: the gene regulatory network of the Caenorhabditis elegans endoderm. Developmental biology. (2002);246(1):68–85. Epub 2002/05/25. doi:10.1006/dbio.2002.0655. PubMed PMID:12027435.

3. Sulston JE, Schierenberg E, White JG, Thomson JN. The embryonic cell lineage of the nematode Caenorhabditis elegans. Developmental biology. (1983);100(1):64–119. Epub 1983/11/01. PubMed PMID:6684600.

4. Lai CH, Chou CY, Ch’ang LY, Liu CS, Lin W. Identification of novel human genes evolutionarily conserved in Caenorhabditis elegans by comparative proteomics. Genome research. (2000);10(5):703–13. Epub 2000/05/16. PubMed PMID:10810093; PubMed Central PMCID:PMCPMC310876.

5. Schubert CM, Lin R, de Vries CJ, Plasterk RH, Priess JR. MEX-5 and MEX-6 function to establish soma/germline asymmetry in early C. elegans embryos. Molecular cell. (2000);5(4):671–82. Epub 2000/07/06. PubMed PMID:10882103.

6. Fraser AG, Kamath RS, Zipperlen P, Martinez-Campos M, Sohrmann M, Ahringer J. Functional genomic analysis of C. elegans chromosome I by systematic RNA interference. Nature. (2000);408(6810):325–30. Epub 2000/12/01. doi:10.1038/35042517. PubMed PMID:11099033.

7. Mello CC, Draper BW, Krause M, Weintraub H, Priess JR. The pie-1 and mex-1 genes and maternal control of blastomere identity in early C. elegans embryos. Cell. (1992);70(1):163–76. Epub 1992/07/10. PubMed PMID:1623520.

8. Ogura K, Kishimoto N, Mitani S, Gengyo-Ando K, Kohara Y. Translational control of maternal glp-1 mRNA by POS-1 and its interacting protein SPN-4 in Caenorhabditis elegans. Development (Cambridge, England). (2003);130(11):2495–503. Epub 2003/04/19. PubMed PMID:12702662.

9. Tsuboi D, Qadota H, Kasuya K, Amano M, Kaibuchi K. Isolation of the interacting molecules with GEX-3 by a novel functional screening. Biochemical and biophysical research communications. (2002);292(3):697–701. Epub 2002/04/02. doi:10.1006/bbrc.2002.6717. PubMed PMID:11922622.

10. Dorfman M, Gomes JE, O’Rourke S, Bowerman B. Using RNA interference to identify specific modifiers of a temperature-sensitive, embryonic-lethal mutation in the Caenorhabditis elegans ubiquitin-like Nedd8 protein modification pathway E1-activating gene rfl-1. Genetics. (2009);182(4):1035–49. Epub 2009/06/17. doi:10.1534/genetics.109.104885. PubMed PMID:19528325; PubMed Central PMCID:PMCPMC2728846.

11. Gomes JE, Encalada SE, Swan KA, Shelton CA, Carter JC, Bowerman B. The maternal gene spn-4 encodes a predicted RRM protein required for mitotic spindle orientation and cell fate patterning in early C. elegans embryos. Development (Cambridge, England). (2001);128(21):4301–14. Epub 2001/10/31. PubMed PMID:11684665.

12. Hallam SJ, Goncharov A, McEwen J, Baran R, Jin Y. SYD-1, a presynaptic protein with PDZ, C2 and rhoGAP-like domains, specifies axon identity in C. elegans. Nature neuroscience. (2002);5(11):1137–46. Epub 2002/10/16. doi:10.1038/nn959. PubMed PMID:12379863.

13. Guedes S, Priess JR. The C. telegans MEX-1 protein is present in germline blastomeres and is a P granule component. Development (Cambridge, England). (1997);124(3):731–9. Epub 1997/02/01. PubMed PMID:9043088.

14. Huang NN, Mootz DE, Walhout AJ, Vidal M, Hunter CP. MEX-3 interacting proteins link cell polarity to asymmetric gene expression in Caenorhabditis elegans. Development (Cambridge, England). (2002);129(3):747–59. Epub 2002/02/07. PubMed PMID:11830574.

15. Pagano JM, Farley BM, McCoig LM, Ryder SP. Molecular basis of RNA recognition by the embryonic polarity determinant MEX-5. The Journal of biological chemistry. (2007);282(12):8883–94. Epub 2007/02/01. doi:10.1074/jbc.M700079200. PubMed PMID:17264081.

16. Sonnichsen B, Koski LB, Walsh A, Marschall P, Neumann B, Brehm M, et al. Full-genome RNAi profiling of early embryogenesis in Caenorhabditis elegans. Nature. (2005);434(7032):462–9. Epub 2005/03/26. doi:10.1038/nature03353. PubMed PMID:15791247.

17. Consortium. CeS. Genome sequence of the nematode C. elegans: a platform for investigating biology. Science (New York, NY). (1998);282(5396):2012–8. Epub 1998/12/16. PubMed PMID:9851916.

18. Bumgarner R. Overview of DNA microarrays: types, applications, and their future. Current protocols in molecular biology. (2013);Chapter 22:Unit 22.1. Epub 2013/01/05. doi:10.1002/0471142727.mb2201s101. PubMed PMID:23288464; PubMed Central PMCID:PMCPMC4011503.

19. Han X, Aslanian A, Yates JR, 3rd. Mass spectrometry for proteomics. Current opinion in chemical biology. (2008);12(5):483–90. Epub 2008/08/23. doi:10.1016/j.cbpa.2008.07.024. PubMed PMID:18718552; PubMed Central PMCID:PMCPMC2642903.

20. Agrawal N, Dasaradhi PV, Mohmmed A, Malhotra P, Bhatnagar RK, Mukherjee SK. RNA interference: biology, mechanism, and applications. Microbiology and molecular biology reviews: MMBR. (2003);67(4):657–85. Epub 2003/12/11. PubMed PMID:14665679; PubMed Central PMCID:PMCPMC309050.

21. Bazan J, Calkosinski I, Gamian A. Phage display--a powerful technique for immunotherapy: 1. Introduction and potential of therapeutic applications. Human vaccines & immunotherapeutics. (2012);8(12):1817–28. Epub 2012/08/22. doi:10.4161/hv.21703. PubMed PMID:22906939; PubMed Central PMCID:PMCPMC3656071.

22. Bruckner A, Polge C, Lentze N, Auerbach D, Schlattner U. Yeast two-hybrid, a powerful tool for systems biology. International journal of molecular sciences. (2009);10(6):2763–88. Epub 2009/07/08. doi:10.3390/ijms10062763. PubMed PMID:19582228; PubMed Central PMCID:PMCPMC2705515.

23. Stein L, Sternberg P, Durbin R, Thierry-Mieg J, Spieth J. WormBase: network access to the genome and biology of Caenorhabditis elegans. Nucleic acids research. (2001);29(1):82–6. Epub 2000/01/11. PubMed PMID:11125056; PubMed Central PMCID:PMCPMC29781.

24. Besser J, Carleton HA, Gerner-Smidt P, Lindsey RL, Trees E. Next-generation sequencing technologies and their application to the study and control of bacterial infections. Clinical microbiology and infection: the official publication of the European Society of Clinical Microbiology and Infectious Diseases. (2018);24(4):335–41. Epub 2017/10/28. doi:10.1016/j.cmi.2017.10.013. PubMed PMID:29074157; PubMed Central PMCID:PMCPMC5857210.

25. Wang Z, Gerstein M, Snyder M. RNA-Seq: a revolutionary tool for transcriptomics. Nature reviews Genetics. (2009);10(1):57–63. Epub 2008/11/19. doi:10.1038/nrg2484. PubMed PMID:19015660; PubMed Central PMCID:PMCPMC2949280.

26. Wang L, Wang S, Li W. RSeQC: quality control of RNA-seq experiments. Bioinformatics (Oxford, England). (2012);28(16):2184–5. Epub 2012/06/30. doi:10.1093/bioinformatics/bts356. PubMed PMID:22743226.

27. Trapnell C, Roberts A, Goff L, Pertea G, Kim D, Kelley DR, et al. Differential gene and transcript expression analysis of RNA-seq experiments with TopHat and Cufflinks. Nature protocols. (2012);7(3):562–78. Epub 2012/03/03. doi:10.1038/nprot.2012.016. PubMed PMID:22383036; PubMed Central PMCID:PMCPMC3334321.

28. Nicolae M, Mangul S, Mandoiu, II, Zelikovsky A. Estimation of alternative splicing isoform frequencies from RNA-Seq data. Algorithms for molecular biology: AMB. (2011);6(1):9. Epub 2011/04/21. doi:10.1186/1748-7188-6-9. PubMed PMID:21504602; PubMed Central PMCID:PMCPMC3107792.

29. Jung N, Bertrand F, Bahram S, Vallat L, Maumy-Bertrand M. Cascade: a R package to study, predict and simulate the diffusion of a signal through a temporal gene network. Bioinformatics (Oxford, England). (2014);30(4):571–3. Epub 2013/12/07. doi:10.1093/bioinformatics/btt705. PubMed PMID:24307703.

30. Szklarczyk D, Franceschini A, Wyder S, Forslund K, Heller D, Huerta-Cepas J, et al. STRING v10: protein-protein interaction networks, integrated over the tree of life. Nucleic acids research. (2015);43(Database issue):D447–52. Epub 2014/10/30. doi:10.1093/nar/gku1003. PubMed PMID:25352553; PubMed Central PMCID:PMCPMC4383874.

31. Stark C, Breitkreutz BJ, Reguly T, Boucher L, Breitkreutz A, Tyers M. BioGRID: a general repository for interaction datasets. Nucleic acids research. (2006);34(Database issue):D535–9. Epub 2005/12/31. doi:10.1093/nar/gkj109. PubMed PMID:16381927; PubMed Central PMCID:PMCPMC1347471.

32. Mitani S. Nematode, an experimental animal in the national BioResource project. Experimental animals. (2009);58(4):351–6. Epub 2009/08/06. PubMed PMID:19654432.

33. Motohashi T, Tabara H, Kohara Y. Protocols for large scale in situ hybridization on C. elegans larvae. WormBook: the online review of C elegans biology. 2006:1–8. Epub 2007/12/01. doi:10.1895/wormbook.1.103.1. PubMed PMID:18050447; PubMed Central PMCID:PMCPMC4781301.

34. Mortazavi A, Williams BA, McCue K, Schaeffer L, Wold B. Mapping and quantifying mammalian transcriptomes by RNA-Seq. Nature methods. (2008);5(7):621–8. Epub 2008/06/03. doi:10.1038/nmeth.1226. PubMed PMID:18516045.

35. Consortium GO. Gene Ontology Consortium: going forward. Nucleic acids research. (2015);43(Database issue):D1049–56. Epub 2014/11/28. doi:10.1093/nar/gku1179. PubMed PMID:25428369; PubMed Central PMCID:PMCPMC4383973.

36. Bateman A, Coin L, Durbin R, Finn RD, Hollich V, Griffiths-Jones S, et al. The Pfam protein families database. Nucleic acids research. (2004);32(Database issue):D138–41. Epub 2003/12/19. doi:10.1093/nar/gkh121. PubMed PMID:14681378; PubMed Central PMCID:PMCPMC308855.

37. Apweiler R, Bairoch A, Wu CH, Barker WC, Boeckmann B, Ferro S, et al. UniProt: the Universal Protein knowledgebase. Nucleic acids research. (2004);32(Database issue):D115–9. Epub 2003/12/19. doi:10.1093/nar/gkh131. PubMed PMID:14681372; PubMed Central PMCID:PMCPMC308865.

38. Shannon P, Markiel A, Ozier O, Baliga NS, Wang JT, Ramage D, et al. Cytoscape: a software environment for integrated models of biomolecular interaction networks. Genome research. (2003);13(11):2498–504. Epub 2003/11/05. doi:10.1101/gr.1239303. PubMed PMID:14597658; PubMed Central PMCID:PMCPMC403769.

39. Mi H, Muruganujan A, Thomas PD. PANTHER in 2013: modeling the evolution of gene function, and other gene attributes, in the context of phylogenetic trees. Nucleic acids research. (2013);41(Database issue):D377–86. Epub 2012/11/30. doi:10.1093/nar/gks1118. PubMed PMID:23193289; PubMed Central PMCID:PMCPMC3531194.

40. Thorpe CJ, Schlesinger A, Carter JC, Bowerman B. Wnt signaling polarizes an early C. elegans blastomere to distinguish endoderm from mesoderm. Cell. (1997);90(4):695–705. Epub 1997/08/22. PubMed PMID:9288749.

41. Abraham T, Pin CL, Watson AJ. Embryo collection induces transient activation of XBP1 arm of the ER stress response while embryo vitrification does not. Molecular human reproduction. (2012);18(5):229–42. Epub 2011/12/14. doi:10.1093/molehr/gar076. PubMed PMID:22155729.

42. Kyriakakis E, Charmpilas N, Tavernarakis N. Differential adiponectin signalling couples ER stress with lipid metabolism to modulate ageing in C. elegans. Scientific reports. (2017);7(1):5115. Epub 2017/07/13. doi:10.1038/s41598-017-05276-2. PubMed PMID:28698593; PubMed Central PMCID:PMCPMC5505976.

43. Desai C, Horvitz HR. Caenorhabditis elegans mutants defective in the functioning of the motor neurons responsible for egg laying. Genetics. (1989);121(4):703–21. Epub 1989/04/01. PubMed PMID:2721931; PubMed Central PMCID:PMCPMC1203655.

44. Kohn RE, Duerr JS, McManus JR, Duke A, Rakow TL, Maruyama H, et al. Expression of multiple UNC-13 proteins in the Caenorhabditis elegans nervous system. Molecular biology of the cell. (2000);11(10):3441–52. Epub 2000/10/12. doi:10.1091/mbc.11.10.3441. PubMed PMID:11029047; PubMed Central PMCID:PMCPMC15005.

45. Pan CL, Baum PD, Gu M, Jorgensen EM, Clark SG, Garriga G. C. elegans AP-2 and retromer control Wnt signaling by regulating mig-14/Wntless. Developmental cell. (2008);14(1):132–9. Epub 2007/12/28. doi:10.1016/j.devcel.2007.12.001. PubMed PMID:18160346; PubMed Central PMCID:PMCPMC2709403.

46. Gottschalk A, Almedom RB, Schedletzky T, Anderson SD, Yates JR, 3rd, Schafer WR. Identification and characterization of novel nicotinic receptor-associated proteins in Caenorhabditis elegans. The EMBO journal. (2005);24(14):2566–78. Epub 2005/07/02. doi:10.1038/sj.emboj.7600741. PubMed PMID:15990870; PubMed Central PMCID:PMCPMC1176467.

47. Yau DM, Yokoyama N, Goshima Y, Siddiqui ZK, Siddiqui SS, Kozasa T. Identification and molecular characterization of the G alpha12-Rho guanine nucleotide exchange factor pathway in Caenorhabditis elegans. Proceedings of the National Academy of Sciences of the United States of America. (2003);100(25):14748–53. Epub 2003/12/06. doi:10.1073/pnas.2533143100. PubMed PMID:14657363; PubMed Central PMCID:PMCPMC299794.

48. Lehner B, Calixto A, Crombie C, Tischler J, Fortunato A, Chalfie M, et al. Loss of LIN-35, the Caenorhabditis elegans ortholog of the tumor suppressor p105Rb, results in enhanced RNA interference. Genome biology. (2006);7(1):R4. Epub 2006/03/02. doi:10.1186/gb-2006-7-1-r4. PubMed PMID:16507136; PubMed Central PMCID:PMCPMC1431716.

49. Bowerman B, Ingram MK, Hunter CP. The maternal par genes and the segregation of cell fate specification activities in early Caenorhabditis elegans embryos. Development (Cambridge, England). (1997);124(19):3815–26. Epub 1997/11/21. PubMed PMID:9367437.

50. Clement A, Solnica-Krezel L, Gould KL. Functional redundancy between Cdc14 phosphatases in zebrafish ciliogenesis. Developmental dynamics: an official publication of the American Association of Anatomists. (2012);241(12):1911–21. Epub 2012/10/03. doi:10.1002/dvdy.23876. PubMed PMID:23027426; PubMed Central PMCID:PMCPMC3508521.

51. Hornsten A, Lieberthal J, Fadia S, Malins R, Ha L, Xu X, et al. APL-1, a Caenorhabditis elegans protein related to the human beta-amyloid precursor protein, is essential for viability. Proceedings of the National Academy of Sciences of the United States of America. (2007);104(6):1971–6. Epub 2007/02/03. doi:10.1073/pnas.0603997104. PubMed PMID:17267616; PubMed Central PMCID:PMCPMC1794273.

52. Bae YK, Barr MM. Sensory roles of neuronal cilia: cilia development, morphogenesis, and function in C. elegans. Frontiers in bioscience: a journal and virtual library. (2008);13:5959–74. Epub 2008/05/30. PubMed PMID:18508635; PubMed Central PMCID:PMCPMC3124812.

53. Hunter CP, Kenyon C. Spatial and temporal controls target pal-1 blastomere-specification activity to a single blastomere lineage in C. elegans embryos. Cell. (1996);87(2):217–26. Epub 1996/10/18. PubMed PMID:8861906.

54. Iioka H, Macara IG. Detection of RNA-Protein Interactions Using Tethered RNA Affinity Capture. Methods in molecular biology (Clifton, NJ). (2015);1316:67–73. Epub 2015/05/15. doi:10.1007/978-1-4939-2730-2_6. PubMed PMID:25967053; PubMed Central PMCID:PMCPMC6047865.

55. Langer-Safer PR, Levine M, Ward DC. Immunological method for mapping genes on Drosophila polytene chromosomes. Proceedings of the National Academy of Sciences of the United States of America. (1982);79(14):4381–5. Epub 1982/07/01. PubMed PMID:6812046; PubMed Central PMCID:PMCPMC346675.

